# Whole genome 6-methyladenosine sequencing (6-mA-Seq) enables bacterial epigenomics studies

**DOI:** 10.1101/2021.02.08.430217

**Authors:** Rachel R. Spurbeck, Angela T. Minard-Smith, Lindsay A. Catlin, Robert W. Murdoch, Rich M. Chou, Kristy Montoya Albrecht

## Abstract

Methylation sequencing using bisulfite treatment has revolutionized the field of molecular biology for eukaryotic systems, unveiling levels of intricacy in regulation of gene expression in response to different environmental conditions. While bacteria also utilize methylation to regulate gene expression, bisulfite sequencing does not work as well in bacteria as in eukaryotes, because bacteria methylate adenosine instead of cytosine. Therefore, global bacterial methylation patterns cannot be studied using common Illumina sequencers. In this work, we demonstrate 6mA-seq, a method that can be used to identify patterns in bacteria that methylate adenosines at GATC sites. Furthermore, this method was used on *Escherichia coli* cultured on four different carbon sources to demonstrate different methylation patterns due to carbon utilization. 6mA-seq can increase the speed in which epigenetic research is conducted in bacteria.

## Introduction

The field of epigenetics has emerged along with next generation sequencing technologies from the study of individual epigenetic marks on specific genes to full genome-wide patterns shifting in response to an environmental stimulus. Epigenetic marks include DNA methylation, histone modifications, and RNA associated silencing, all which control gene or protein expression. Epigenetics is most studied in multicellular organisms such as humans and plants, where epigenetic marks function in embryogenesis, cellular differentiation, genomic imprinting, and play roles in pathogenesis of diseases such as cancer [1]. Most techniques for the study of epigenetics were developed to understand the epigenomes of eukaryotes and are not tuned specifically to account for a set of additional marks utilized by prokaryotes. Thus, new tools are required to facilitate and determine microbial epigenomes.

While the set of epigenetic marks is smaller in bacteria than in eukaryotes, as microbes do not have histones, epigenetics still regulate gene and protein expression. Beyond transcription factors, microbes deploy small RNAs and DNA methylation to regulate gene and protein expression. Small RNAs can readily be studied using commercial-available small RNA sequencing kits. Few techniques are available to detect the most common methylation mark in bacteria, 6-methyladenosine (6mA) [1], and those require less common sequencing equipment such as PacBio [1, 2] or Oxford Nanopore Technologies [3] sequencing platforms. Global methylation patterns currently are analyzed using bisulfite sequencing which takes advantage of the chemical transformation of unmethylated cytosine into uracil by treatment with bisulfite. Methylation of cytosine protects the cytosine from transformation, and therefore, when compared to a known genome sequence, one can deduce which bases were methylated as the bases that remained cytosine after bisulfite treatment. This method can be utilized on the most common sequencing platform from Illumina. However, bisulfite sequencing does not detect adenosine methylation. While both marks are present in the genomes of eukaryotes and prokaryotes [4, 5], higher eukaryotes mainly utilize 5-methylcytosine (5mC), while bacteria commonly utilize 6mA [1, 6].

The role of DNA methylation in bacteria ranges from defense against bacteriophage through restriction-modification systems, initiation of DNA replication, DNA repair, and gene regulation [5, 7–10]. A tool that could enable global methylation pattern analysis of 6mA on a common sequencing platform (e.g., Illumina DNA sequencing platforms) would expand 16S sequencing and metagenomics to include valuable epigenetics research. [11]. Furthermore, there is direct evidence linking 6mA to several species of eubacteria, mosquitos, wheat, protists, and indirect evidence links 6mA to several archaea and vertebrates [12, 13], raising the possible utility of a 6mA tool beyond microbial characterization. Presently, the only method for detecting global 6mA patterns is to use PacBio sequencing or MinION sequencing. These instruments are not as widely distributed as Illumina sequencers and are known to have higher error rates [11], therefore a 6mA-sequencing solution on Illumina instruments would increase the quality and reliability of experimental results. PacBio has also been shown to overestimate 6mA, especially in eukaryotic genomes where the modification is rare [14]. By enabling 6mA to be detected on Illumina sequencers, microbial epigenomics can take advantage of a platform with the highest base calling accuracy. In this work, a method to sequence 6mA patterns on an Illumina sequencer was developed and used to characterize *Escherichia coli* global 6mA pattern shifts when cultured on different carbon sources.

Adenine methylation in bacteria is controlled by two types of methyl transferases: those that methylate upon DNA replication to enable restriction modification systems to distinguish self from non-self, and methyl transferases that respond to environmental changes effecting transcription [5, 7]. Methyl transferases used to distinguish self from non-self produce stable patterns, so are not useful in identifying environmental effects on microbes. However, 6mA is produced by the action of at least two methyl transferases not associated with self-recognition: DAM methylase, which methylates at GATC sites, and CcrM, which methylates at the motif GANTC [7]. The method demonstrated herein exploits these modification pathway differences by using restriction endonucleases that cut DNA at GATC or GANTC that are sensitive to adenine methylation. When 6mA is present, the restriction enzymes cannot cut GATC or GANTC. Therefore, like bisulfite sequencing, one can look for methylated bases by looking for places in the genome not affected by the enzymatic treatment. This method was tested on two *E. coli*: one that had a mutation in DNA adenine methylase (DAM), and so could not methylate its DNA, and a wild-type, methylation competent strain. This showed 6mA-seq is highly sensitive to methylation. Subsequently, the methylation competent *E. coli* was cultivated under different carbon conditions and analyzed by 6mA-seq to identify methylation pattern changes. This study has demonstrated that methylation changes occur at sites in proximity to genes known to be involved in metabolism, thus highlighting that methylation sequencing in bacteria identifies biologically relevant epigenetic patterns.

## Materials and Methods

### Bacterial strains and culture conditions

Two *Escherichia coli* strains were cultured in 10 mL of Luria broth overnight at 37°C with aeration. *E. coli* ATCC25922 has a wildtype genotype with both *dam* and *dcm* activity, enabling it to methylate its genome at both cytosine and adenine. The second strain was a *dam*^-^/*dcm*^-^, derivative of *E. coli K-12*, not capable of methylation at GATC or cytosine sites. Both strains harbored mutations that effect methylation at GANTC sites.

### Sample preparation and sequencing

DNA was extracted in triplicate from both *E. coli* cultures using the DNeasy UltraClean Microbial Kit from Qiagen according to the manufacturer’s instructions. The triplicates were then pooled for each strain and then quantified by Quantus to calculate the correct volumes for optimal input into the library preparation for each sample. Each sample was diluted 1:10 to enable fluorimetry readings to fall within the standard curve using the Quantus Fluorometer ONE dsDNA protocol which has an upper limit of 400 ng/μL. 1 μg of DNA from each sample was then aliquoted and brought up to 20 μL with sterile water for a total of three aliquots per strain. Each 20 μL aliquot was then fragmented by Covaris S220 using the MicroTUBE-15 AFA Bead protocol for 300 bp fragmentation. Sequencing libraries were prepared using the Kapa Hyper kit with amplification module and indexed using unique single-end indices for each library with the following modifications from the manufacturer’s protocol. After index ligation, the libraries went through one 0.8x ratio bead clean up to remove extra reagents and complete a buffer exchange into Tris buffer. The libraries were then restriction digested with MboI, DpnII, or HinFI. The six library preparations then went through another bead clean up and then amplified for 15 cycles and bead cleaned according to the manufacturer’s instructions. The libraries were quantified by Qubit and visualized by Agilent Bioanalyzer. The libraries were normalized to 2 nM and pooled for 300 bp paired-end sequencing on the Illumina MiSeq with a MiSeq V2 300 cycle cartridge.

### Demonstration of utility in methylation profiling

*E. coli* ATCC25922 (*dam*^+^/*dcm*^+^) cultured for isolated colonies on LB agar. A single colony was used to inoculate one 10 mL Luria broth culture which incubated overnight with aeration. This culture was then used to inoculate triplicate subcultures in Luria broth, M9-glucose, M9-glycerol, or M9 sorbitol overnight at 37°C with aeration. DNA was extracted from the 12 subcultures using the DNeasy UltraClean Microbial Kit from Qiagen according to the manufacturer’s instructions and quantified by Quantus. Each sample was diluted 1:10 to enable fluorimetry readings to fall within the standard curve using the Quantus Fluorometer ONE dsDNA protocol which has an upper limit of 400 ng/μL. 20 μL aliquots at a concentration of 50 ng/μL were then fragmented by Covaris S220 using the MicroTUBE-15 AFA Bead protocol for 300 bp fragmentation. Sequencing libraries were prepared using the Kapa Hyper kit and indexed using unique single-end indices for each library with the following modifications from the manufacturer’s protocol. After index ligation, the libraries went through one 0.8x ratio bead clean up to remove extra reagents and complete a buffer exchange into Tris buffer. The libraries were then restriction digested with DpnII for 60 minutes. The libraries went through another bead clean up and then amplified for 15 cycles and bead cleaned according to the Kapa Hyper with amplification module instructions. The libraries were quantified by Qubit and visualized by Agilent Bioanalyzer. The libraries were normalized to 2 nM and pooled for 300 bp paired-end sequencing on the Illumina MiSeq with a MiSeq V2 300 cycle cartridge.

### Bioinformatics

Raw fastq data were trimmed using Cutadapt v.1.16. Cutadapt trimmed off low quality ends with a quality lower than Q20 and removed read pairs that were less than 200bp in either read. The fastx toolkit (Hannon lab) fastx_quality_stats created quality data tables for the raw reads and the fastx toolkit fastq_quality_boxplot_graph.sh script produced quality plots for each sample and read direction that were used to set the quality cutoff in Cutadapt.

The filtered and trimmed reads were aligned by BWA v.0.7.156 respectively to the matching genome: either *E. coli* K12 (*dam*^-^/*dcm*^-^ samples) or to the *E. coli* ATCC25922 genome (*dam*^+^/*dcm*^+^) samples. The alignments were visualized in IGV and an R script was used with the Rsamtools package to generate depth of coverage for each alignment. Using the IGV tool ‘Find Motif’ the nucleotide coordinates for every location where the sequence motifs GATC and GANTC were present in each *E. coli* genome was identified and a BED file was exported for each motif. The sequence depth of coverage over the GATC or GANTC motifs was then calculated using BEDtools coverageBed. Coverage of GATC sites were normalized to the full genome coverage. Normalized GATC sites were then determined to be either hypomethylated or hypermethylated based on the relative coverage of the GATC sites compared to the full genome coverage. Sites that were at least 2 standard deviations above the average coverage were considered hypermethylated and those at least 2 standard deviations below the average coverage were considered hypomethylated.

## Results

### 6mA-Seq development using methylation negative and positive *E. coli* strains

This study was conducted using two strains of *Escherichia coli* to determine if the 6-methyl adenine sequencing method (6mA-Seq, Fig. 1) was feasible. The selected strains represent an unmethylated and a methylated strain at GATC sites. Three enzymes were chosen for method development: MboI, DpnII, and HinFI. MboI and DpnII both have endonuclease activity and cut 5’ of the G at the GATC motif. Both enzymes cannot cut when the DNA is methylated. HinFI cuts between the G and A in GANTC motifs. GANTC is the motif methylated by CcrM in alpha proteobacteria for regulation of the cell cycle. *E. coli* does not have *ccrM*, and therefore cannot be protected by methylation at the adenine in the motif but this enzyme can be blocked by some combinations of overlapping CpG methylation. For the *dam*^-^/*dcm*^-^ strain GATC motif, no protection was expected following treatment with MboI or DpnII, with similar outcomes expected for the GANTC motif when treated by HinFI. Conversely, for the *dam^+^/dcm^+^* strain, it was expected that most GATC sites would be protected from digestion and that there could be some low-level protection from HinFI at sites that may have CpG methylation.

**Fig. 1.**
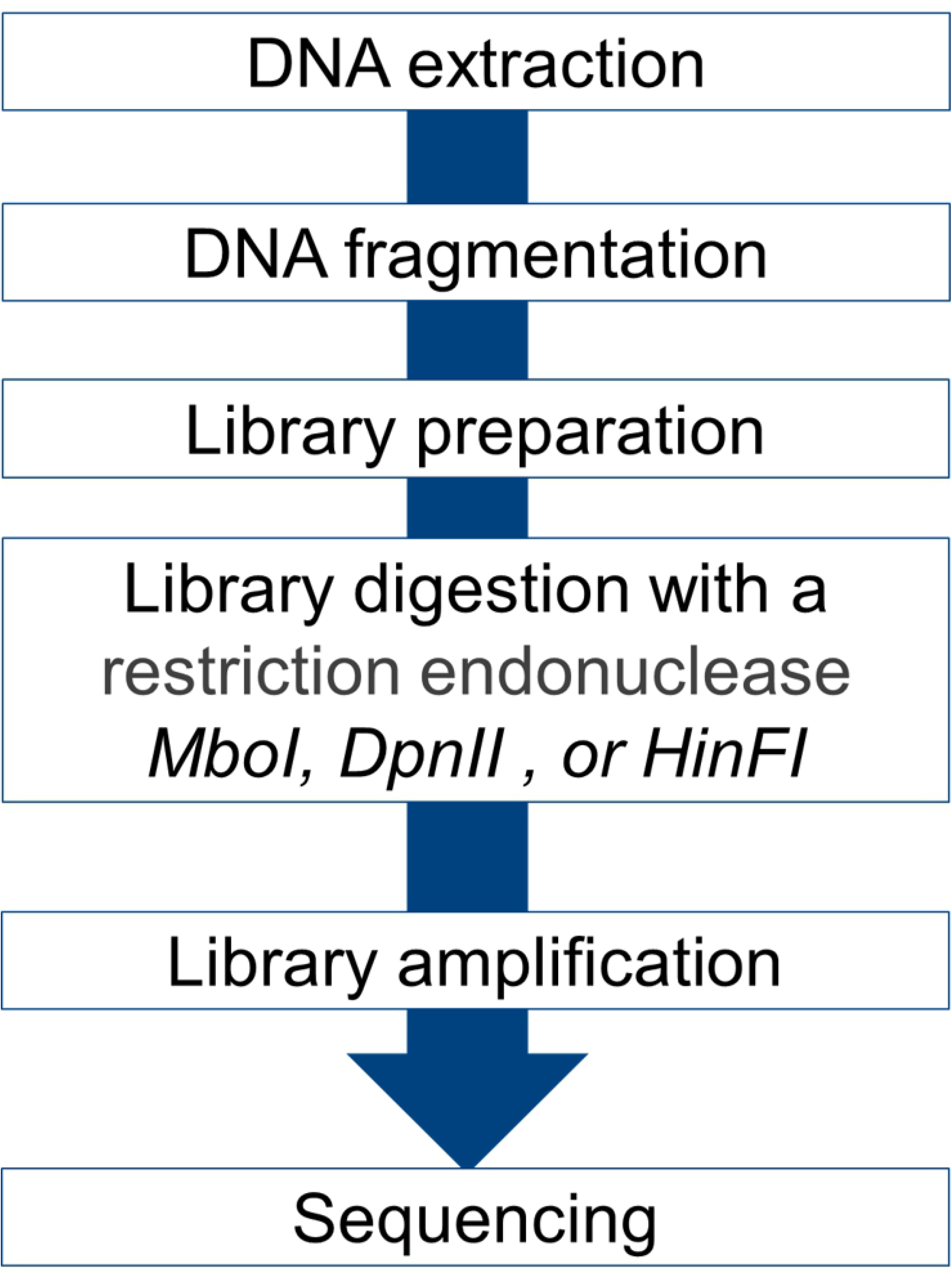
6mA-Seq workflow.

PCR-free Illumina sequencing libraries were prepared from DNA from both strains. These libraries were then treated with the restriction enzymes that removed library molecules that contain unmethylated adenines at the respective motifs leaving behind only molecules at potential 6mA sites that were methylated (Fig. 2). The *dam*^-^/*dcm*^-^ strain had 19146 possible 6mA GATC sites of which none were methylated according to the 6mA sequencing analysis, whereas the *dam*^+^/*dcm*^+^ strain had 98.8% of its 20460 possible 6mA GATC sites methylated (Table 1). As a control, the number of GATC sites that had coverage was calculated for HinFI treated samples, which cuts at GANTC, not GATC sites. In the *dam*^-^/*dcm*^-^ strain treated with HinFI, there was a total genome coverage of 115.7x, similar to that of the *dam*^+^/*dcm*^+^ strain which had a total genome coverage of 104.7x. Both strains had coverage at 97.8% of GATC sites when treated with HinFI. These results show the utility of our method for detection of differential 6mA. Assuming that 6mA occurs at similar locations in all species, this method should be expandable to detect differential 6mA methylomes across the tree of life.

**Figure 2.**
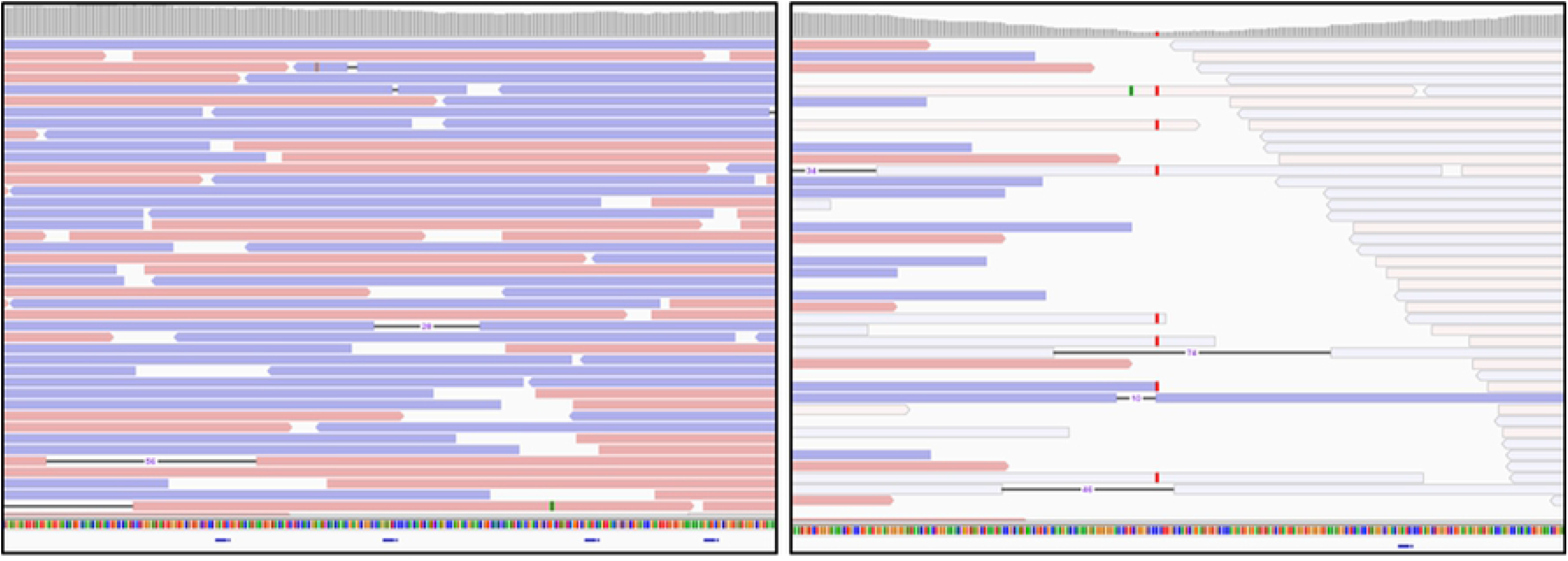
IGV images of regions containing potential 6mA sites. The blue line at bottom of images indicates GATC sites. The left image depicts four 6mA sites in the *dam*^+^/*dcm*^+^ *E. coli* strain that were methylated and the right image depicts one predicted 6mA site that was not methylated, and therefore was not covered by sequence.

**Table 1.**
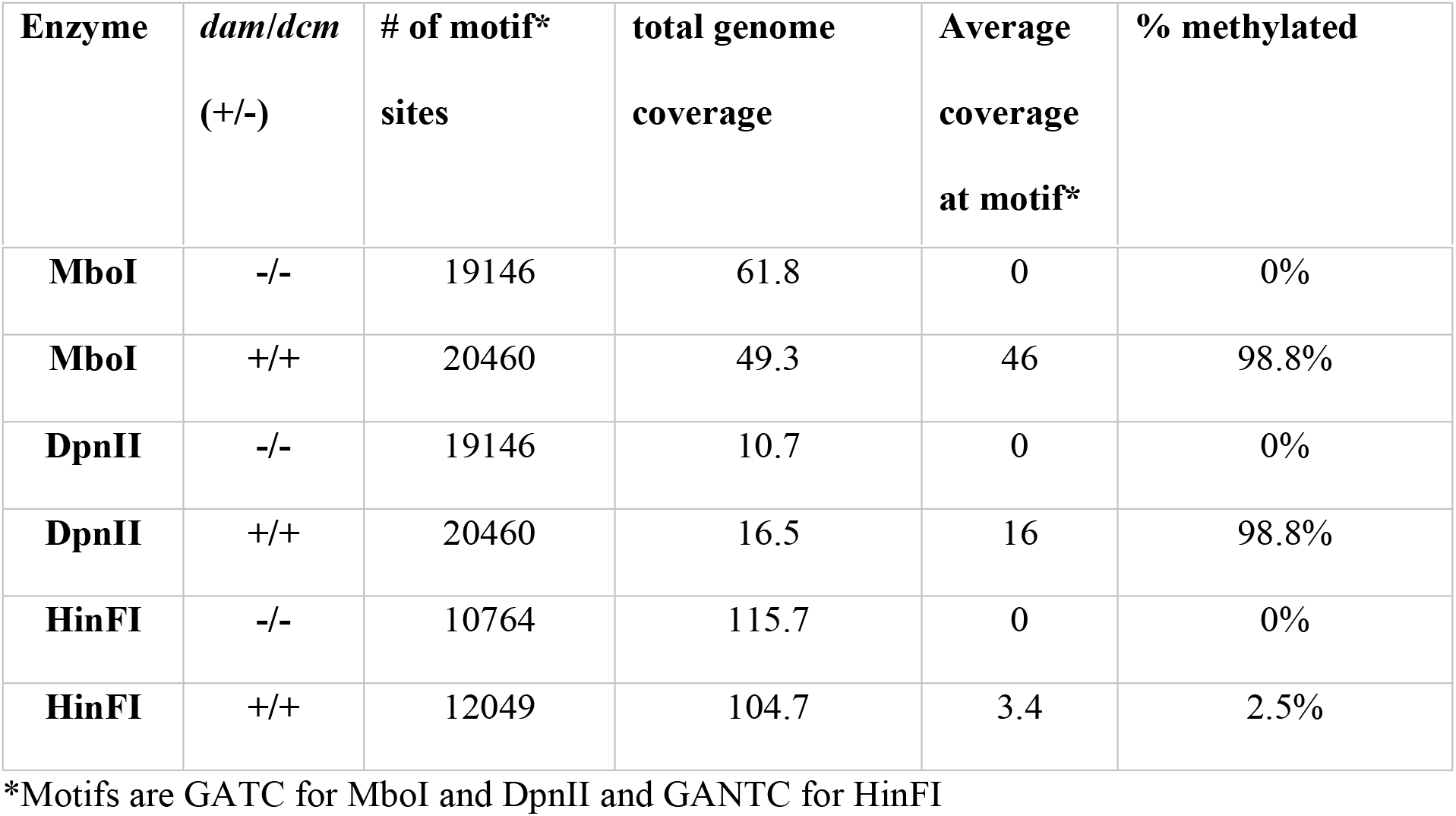
6mA sequencing coverage statistics for *dam*^-^/*dcm*^-^ and *dam*^+^/*dcm*^+^ *E. coli* strains.

### Methylation profiles shift due to carbon source in methylation competent *E. coli*

To determine if the 6mA-seq method could be used to identify methylation profile changes in methylation competent bacteria, *E. coli* ATCC25922 was cultured in four different carbon source conditions and the methylation profile identified for triplicate cultures. The average genome coverage was calculated for each library and the average coverage across GATC sites identified. Hyper and hypo methylated sites were identified as those with higher or lower than two standard deviations from the mean coverage and consistent across the triplicate cultures. Culture media were Luria-Bertani broth or M9 minimal salts medium. Three different carbon sources were compared in minimal media: glycose (M9-glucose), glycerol (M9-glycerol) and sorbitol (M9-sorbitol). *E. coli* ATCC25922 was incubated with shaking overnight in each liquid media overnight at 37°C 5% CO_2_. A total of 220 GATC sites demonstrated hyper or hypomethylation in *E. coli* ATCC25922 (Fig. 3). Three GATC sites (position start: 777328, 3928044, and 5032297, corresponding to genes encoding transcriptional regulator *gutM*, Poly-beta-1,6-N-acetyl-D-glucosamine N-deacetylase *pgaB*, and Salicyl-AMP ligase *ybtE*) were consistently hypomethylated and two (position start: 2907398 and 2907498, both within the gene encoding LPS-assembling protein, *lptD*) were consistently hypermethylated in all four conditions suggesting that the methylation at these sites was not influenced by carbon source. 8 GATC sites (position start: 1661619, 2581408, 2581441, 2581801, 2582072, 2582134, 2718715, 2906872, and 2907142) were consistently hypermethylated in M9 minimal media regardless of carbon source and were not in LB. 9 GATC sites were hypomethylated only in LB (position start: 409277, 479355, 1579192, 1684645, 2181531, 2498588, 2957634, 3782914, and 4031299). LB shared 6 hypomethylated GATC sites with M9-glycerol and M9-glucose (position start: 665850, 786885, 3782985, 3928100, 5031607, and 5032236). Of the hypermethylated GATC sites, 54 were in intragenic regions and 105 were intragenic. Similarly, 15 hypomethylated GATC sites were present at intergenic regions and 46 were intragenic.

**Figure 3.**
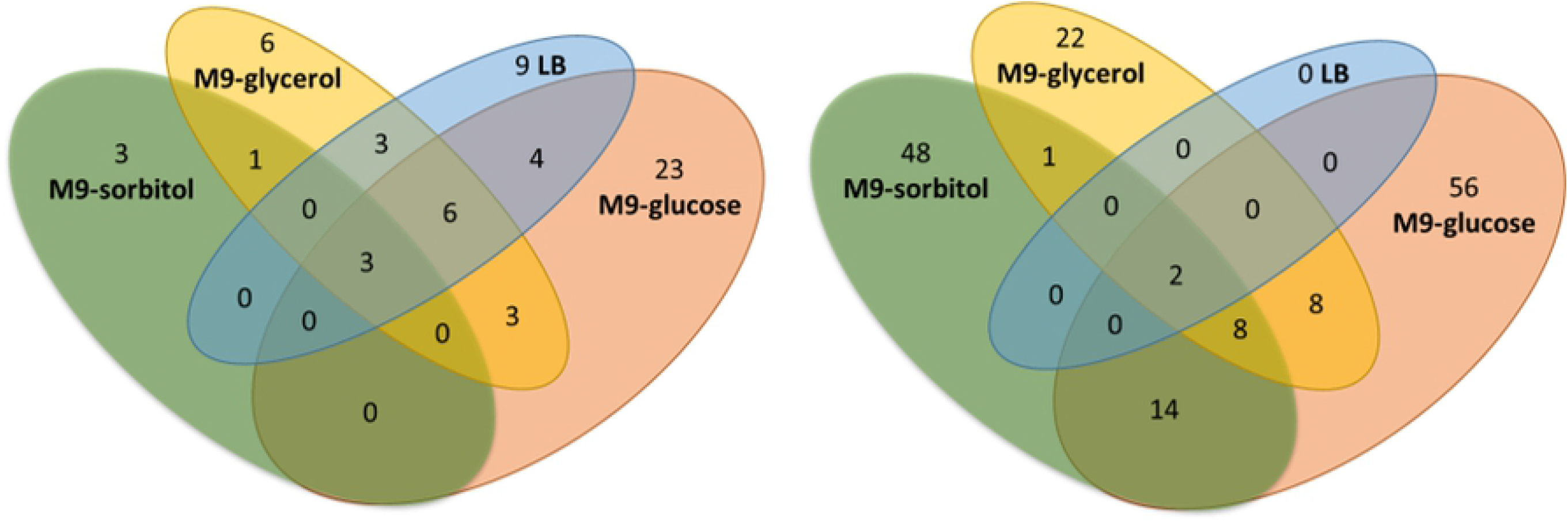
Venn diagrams of A. hypomethylated GATC sites and B. hypermethylated GATC sites in *E. coli ATCC25922* dependent on carbon source.

### Functional analysis of hyper and hypomethylated regions

To begin to understand the potential function of methylation in transcriptional control due to carbon source, the loci identified as hyper or hypomethylated were annotated and sorted by gene function (Fig. 4). In all culture conditions, the majority of GATC sites identified were in regions of unidentified function. However, some functional groups were differentially methylated based on carbon source, including carbohydrate metabolism, and lipopolysaccharide synthesis (Fig. 4).

**Figure 4.**
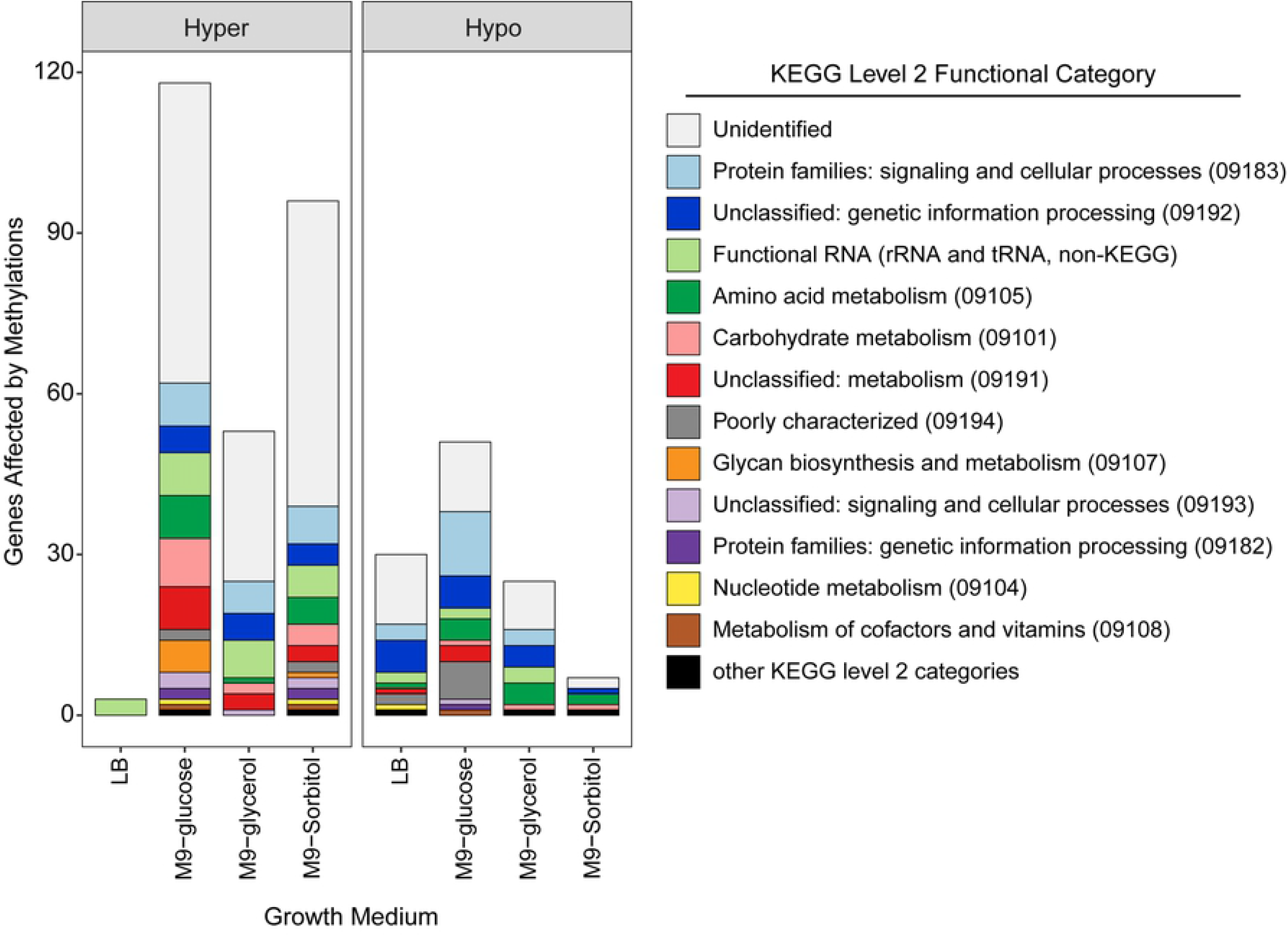
KEGG orthology level 2 functional categories of genes affected by methylation. Sites were identified within the coding region (intragenic) or at an adjacent site (intergenic) and determined to be hyper or hypomethylated. Functional RNA encoding genes (rRNA and tRNA) were predicted using barrnap v0.9-3.

Of the regions in which functional annotations could be made, fifteen hypermethylated loci were present in genes related to carbohydrate metabolism, with most methylation occurring when cultured in glucose, followed by sorbitol and glycerol, and three genes were hypomethylated. No carbohydrate metabolism genes were hyper- or hypomethylated when cultured in LB. In M9-sorbitol, two intragenic GATC sites in were hypermethylated in pectate disaccharide-lyase (*pelX*), involved in pentose metabolism. Two intergenic GATC sites were also hypermethylated in M9 sorbitol upstream of genes encoding enzymes involved in central carbon metabolism systems such as TCA and pyruvate metabolism: aconitate hydratase (*acnA*) 2-isopopylmalate synthase (*leuA*).

Culture in M9-glucose led to the most carbohydrate-related methylated GATC sites (9), most of which specifically involved sugar metabolism. Four methylated GATC sites were in or adjacent to genes encoding enzymes involved in sugar phosphorylation or metabolism of phosphosaccharides including fructose (*fruA*), sorbose (*sorB*), and glycogen (*glgP*). One methylated GATC was in glucose-galactose epimerase (*galE*). Methylated GATC sites were also found in phospho-beta-glucosidase (*bglE*), likely involved in starch synthesis from glucose-6-phosphate; mannonate dehydratase (*manD*), possibly involved in channeling glucose to pentose sugars; alcohol dehydrogenase, a key enzyme for fermentation of glucose; and 2-isopropylmalate synthase (*leuA*), involved in central pyruvate metabolism. As mentioned above, *leuA* was also hypermethylated when grown on sorbitol at the same GATC site. *E. coli* cultured in M9-glycerol had two hypermethylated GATC sites involved in carbohydrate metabolism: one intragenic GATC site in oligogalacturonide lyase (*ogl*), which is involved in pentose and glucuronate interconversions, and one GATC site upstream of galactarate dehydratase (*garD*).

Each carbon source in M9 minimal media led to one substrate-specific intragenic methylation, none of which were present in the hypermethylation group. All three were in genes related to pyruvate metabolism: Sorbitol: phosphoenylpyruvate carboxylase *ppc*, Glucose: malate dehydrogenase *maeA*, and Glycerol: pyruvate-ferredoxin/flavodoxin oxidoreductase *nifJ*.

Culture in M9-glucose lead to several intragenic hypermethylated GATC sites in genes related to lipopolysaccharide synthesis, including *waaR, waaV, waaL*, two in *waaY*, and *eptC*. In contrast, the only other instance of a methylation of the lipopolysaccharde system gene was of *eptC* in when hypermethylation group was grown in sorbitol.

## Discussion

This work has demonstrated the potential utility of 6mA-Seq to detection of differentially methylated loci in bacteria. A *dam*^-^ mutant *E. coli* strain was used to demonstrate 0% of GATC sites protected against restriction digestion as compared to 98.8% of GATC sites protected in the wild-type *E. coli*. Using 6mA-Seq were we able to demonstrate methylation pattern shifts due to carbon utilization across the *E. coli* genome. The methylation pattern of *E. coli* cultured in LB, a nutrient rich medium demonstrated significantly less hyper- and hypomethylated sites suggesting that *E. coli* was less restricted in this environment, and able to express genes to utilize multiple nutrients present in the medium. In contrast, under minimal medium conditions, several genes were hyper or hypomethylated demonstrating a shift in the genetic content available for expression due to resource limitation, with only three shared hypomethylated sites (biosynthetic threonine ammonia-lyase, a tRNA-Ala and a location ~100 bp from this tRNA-Ala) and two shared hypermethylated (IS3 family transposase and ClbS/DfsB family four-helix bundle protein) in all three carbon sources. Furthermore, each carbon source contributed to unique methylation patterns with different GATC sites being hyper or hypomethylated in carbon metabolism and lipopolysaccharide biosynthesis pathways. Future studies will be necessary to determine if the methylation patterns identified correlate with gene expression patterns in the same growth conditions, as methylation at promoters or intragenic regions in bacteria can increase, decrease, or have no effect on gene transcription [15], due to the complex interactions between methylation and transcription factors.

This new 6mA-seq method enables researchers to pursue epigenetics of bacteria and lower eukaryotes at a base that cannot be queried by traditional methods. Several restriction enzymes are highly sensitive to methylation, and by taking advantage of this sensitivity new methylation patterns can emerge across the genome. However, this method is limited in that it only enables identification of methylation at the restriction enzyme motifs and requires ~30x sequencing coverage for accuracy and sensitivity, which becomes significantly more expensive as one moves to larger genome sizes such as higher eukaryotes, where 6mA has been discovered at extremely low levels (0.00006-0.00077% in human, rat, and plants) [13]. Furthermore, 6mA methylation motifs may be organism specific [16], suggesting that 6mA-seq will need to be expanded to include a cocktail of enzymes to detect the epigenetic diversity in nature.

## Acknowledgements

This work was funded by internal research and development funding from Battelle Memorial Institute.

## Supporting information captions

**Supporting Information Data Table 1.** Genome sequence coverage at all motif sites for *dam*^-^/*dcm*^-^ and *dam*^+^/*dcm*^+^ *E. coli* treated with different methylation sensitive restriction enzymes.

**Supporting Information Data Table 2.** Normalized genome sequence coverage at all motif sites over two standard deviations from the mean genome coverage for *E. coli* cultured in LB, M9-sorbitol (M9S), M9-glucose (M9glu), and M9-glycerol (M9gly) after 6mAseq.

